# Histone Deacetylase 11 is an ε-*N*-Myristoyllysine Hydrolase

**DOI:** 10.1101/211839

**Authors:** Carlos Moreno-Yruela, Iacopo Galleano, Andreas S. Madsen, Christian A. Olsen

## Abstract

Histone deacetylase (HDAC) enzymes are important regulators of diverse biological function, including gene expression, rendering them potential targets for intervention in a number of diseases, with a handful of compounds approved for treatment of certain hematologic cancers. Among the human zinc-dependent HDACs, the most recently discovered member, HDAC11, is the only member assigned to subclass IV, the smallest protein, and the least well understood with regards to biological function. Here we show that HDAC11 cleaves long chain acyl modifications on lysine side chains with remarkable efficiency compared to acetyl groups. We further show that several common types of HDAC inhibitors, including the approved drugs romidepsin and vorinostat, do not inhibit this enzymatic activity. Macrocyclic hydroxamic acid-containing peptides, on the other hand, potently inhibit HDAC11 demyristoylation activity. These findings should be taken carefully into consideration in future investigations of the biological function of HDAC11 and will serve as a foundation for the development of selective chemical probes targeting HDAC11.

## INTRODUCTION

Lysine deacylases (KDACs) are hydrolases conserved across archaea, bacteria, and eukaryotes, which cleave posttranslational acylation of the ε-*N*-amino groups of lysine residues in the proteome. Humans have a total of eighteen KDACs; the eleven zinc-dependent HDACs, which are categorized in class I (HDACs 1–3 and 8), class IIa (HDACs 4, 5, 7, and 9), class IIb (HDACs 6 and 10), and class IV (HDAC11) (Fig. 1A) (Marmorstein, 2001; Gregoretti et al., 2004). Class III KDACs comprise the structurally and mechanistically distinct NAD^+^-dependent sirtuins (SIRT1–7) (Frye, 2000). Among the eleven zinc-dependent enzymes, class I and IV enzymes are primarily located in the nucleus, class IIa enzymes are shuttling between nucleus and cytosol, and class IIb enzymes are mainly localized to the cytosol (de Ruijter et al., 2003). In addition to their initially discovered epigenetic effects achieved through regulation of ε-*N*-acetyllysine (Kac) levels in histones, HDACs have been implicated in diverse biology including metabolism, inflammation, circadian rhythm, as well as neurological development and brain function (Guan et al., 2009; Villagra et al., 2009; Gräff et al., 2012; Jakovcevski and Akbarian, 2012; Falkenberg and Johnstone, 2014; Tough et al., 2016). Recently, it has become evident that lysine residues may be decorated with a variety of acyl groups other than acetyl, and certain sirtuins have been shown to target these posttranslational modifications (PTMs) (Bheda et al., 2016). For example, SIRT4 has been shown to cleave ε-*N*-lipoyllysine (Klip) (Mathias et al., 2014) and ε-*N*-(3-methylglutaconyl)lysine (Kmgc) (Anderson et al., 2017), SIRT5 regulates ε-*N*-malonyllysine (Kmal) (Du et al., 2011; Peng et al., 2011; Hirschey and Zhao, 2015; Nishida et al., 2015), ε-*N*-succinyllysine (Ksuc) (Du et al., 2011; Park et al., 2013; Rardin et al., 2013; Hirschey and Zhao, 2015), and ε-*N*-glutaryllysine (Kglu) levels (Tan et al., 2014; Hirschey and Zhao, 2015), while SIRT2 (Feldman et al., 2013; Galleano et al., 2016; Madsen et al., 2016) and SIRT6 (Jiang et al., 2013; Madsen et al., 2016) efficiently cleave ε-*N*-myristoyllysine (Kmyr). For the zinc-dependent HDACs, on the other hand, only few examples of deacylation activities other than deacetylation have been reported and these have been weaker than their potencies against Kac (Madsen and Olsen, 2012a; Aramsangtienchai et al., 2016; Wei et al., 2017). Phylogenetically, the human zinc-dependent HDACs appear in three different clusters, which has given rise to their classification (Fig. 1A). The primary targets of class I isozymes are Kac residues in histones and the class IIb isozyme, HDAC6, has been associated with deacetylation of several cytosolic substrates including α-tubulin and p53 (Hubbert et al., 2002; Zhang et al., 2003; Ryu et al., 2017). However, the remaining HDACs, albeit involved in regulation of diverse biological mechanisms (Verdin et al., 2003; Chen and Cepko, 2009; Villagra et al., 2009), have been difficult to pair with specific protein targets and several HDACs, i.e. class IIa isoforms 4, 5, 7, and 9 as well as HDACs 10 and 11, exhibit low deacetylase activity in vitro. As found for the sirtuins, where several isozymes have been shown to preferentially cleave other ε-*N*-acyllysine modifications than Kac, we therefore envisioned that this might also be the case for members of the zinc-dependent HDAC isozyme classes. Here, we present the screening for activity of all eleven HDACs against a diverse selection of ε-*N*-acyllysine substrates. Surprisingly, we found HDAC11 to exhibit robust activity against long chain acyl groups and disclose kinetic evaluation of this novel enzymatic activity as well as inhibition thereof using macrocyclic HDAC inhibitors, which should be of great importance for the continued efforts to elucidate the biology of HDAC11.

**Figure 1.**
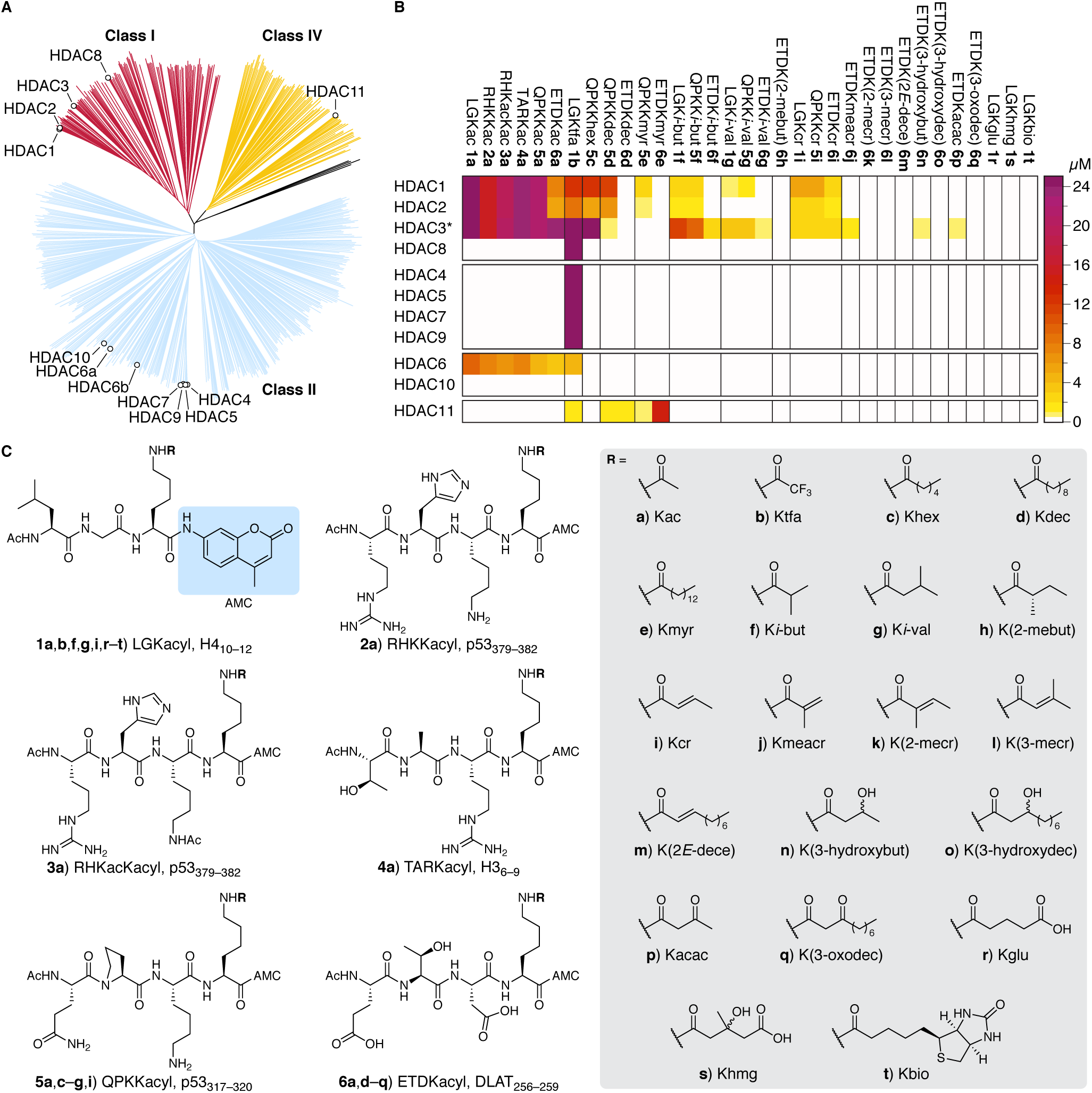
Profiling of the activities of HDAC1–11 against a series of 33 fluorescent substrates. (A) Phylogenetic representation of 602 HDACs in the NCBI Conserved Protein Domain HDAC-family (cd09301), highlighting classes I, II, and IV and labeling the human HDACs. (B) Heat map showing the activities of recombinant HDAC1–11 against the full substrate series. Please consult Figure S1 for bar graph representation of the data. The results are based on at least two individual end-point assays performed in duplicate. (C) Structures of the screened fluorophore-coupled substrates. AMC = 7-amino-4-methylcoumarin.

## RESULTS

### Substrate screening

For the initial screening against an in-house library of more than thirty fluorogenic substrates (**1a**–**6q**), we applied all eleven zinc-dependent human HDACs using conditions described for commercially available assay kits (Fig. 1B and 1C). The substrate series included commercially available substrates recommended for certain HDACs (**2a**, **3a**, and **5a**) and a diverse series of acyl groups, some of which were synthesized on several scaffolds. The acyl groups were chosen to cover PTMs demonstrated to exist by MS/MS proteomics-based methods as well as additional potential modifications that in principle could arise from lysine residues reacting with the corresponding acyl-CoA species.

Not surprisingly, we recorded robust deacetylation activity of HDACs 1–3 and 6 (Fig. 1B) and moderate activity of HDAC8 (Supplemental Fig. S1), which may require the nucleosome structure to exhibit potent activity against Kac. As previously reported, class IIa isozymes (4, 5, 7, and 9) as well as HDAC8 exhibited highly potent activities against the non-physiological ε-*N*-trifluoroacetyllysine (Ktfa) substrate (Riester et al., 2004; Bradner et al., 2010). The second class IIb enzyme, HDAC10, did not show significant conversion of any of our substrates, which is in agreement with a recent report demonstrating the ability of HDAC10 to deacylate polyamines rather than lysine residues (Hai et al., 2017). Interestingly, however, HDAC11 cleaved Kmyr with high efficiency equal to the conversions of Kac recorded for class I HDACs (orange color in the heat map, Fig. 1B). This was a surprising finding that we decided to investigate in further detail because of its potential impact on the currently limited understanding of the function of HDAC11.

### Effects of buffer conditions, HDAC11 construct, and substrate structure

In previous investigations of the demyristoylase activity of SIRT2, we encountered differences in enzyme efficiency depending on the applied buffer. Moreover, the commercial Tris buffer is at pH 8, which is not physiologically relevant for cytosolic and nuclear proteins, prompting us to test the HEPES buffer (pH 7.4) described by Bradner and Mazischek (Bradner et al., 2010). Finally, due to the potential of sequestering fatty acid-containing substrate by bovine serum albumin (BSA) in the buffer, we included a modified version of the latter containing only 0.05 mg/mL BSA. In addition to the buffer, we compared the activities of two commercially available HDAC11 constructs [one untagged (light blue bars) and one GST-tagged (red bars)] at varying concentrations in all three buffers (Fig. 2A). While the buffer showed a minor effect except at sub-nanomolar enzyme concentrations, the GST-tag appeared to cause a striking decrease in activity (Fig. 2A) and we decided to continue by using the untagged HDAC11 for further investigations.

**Figure 2.**
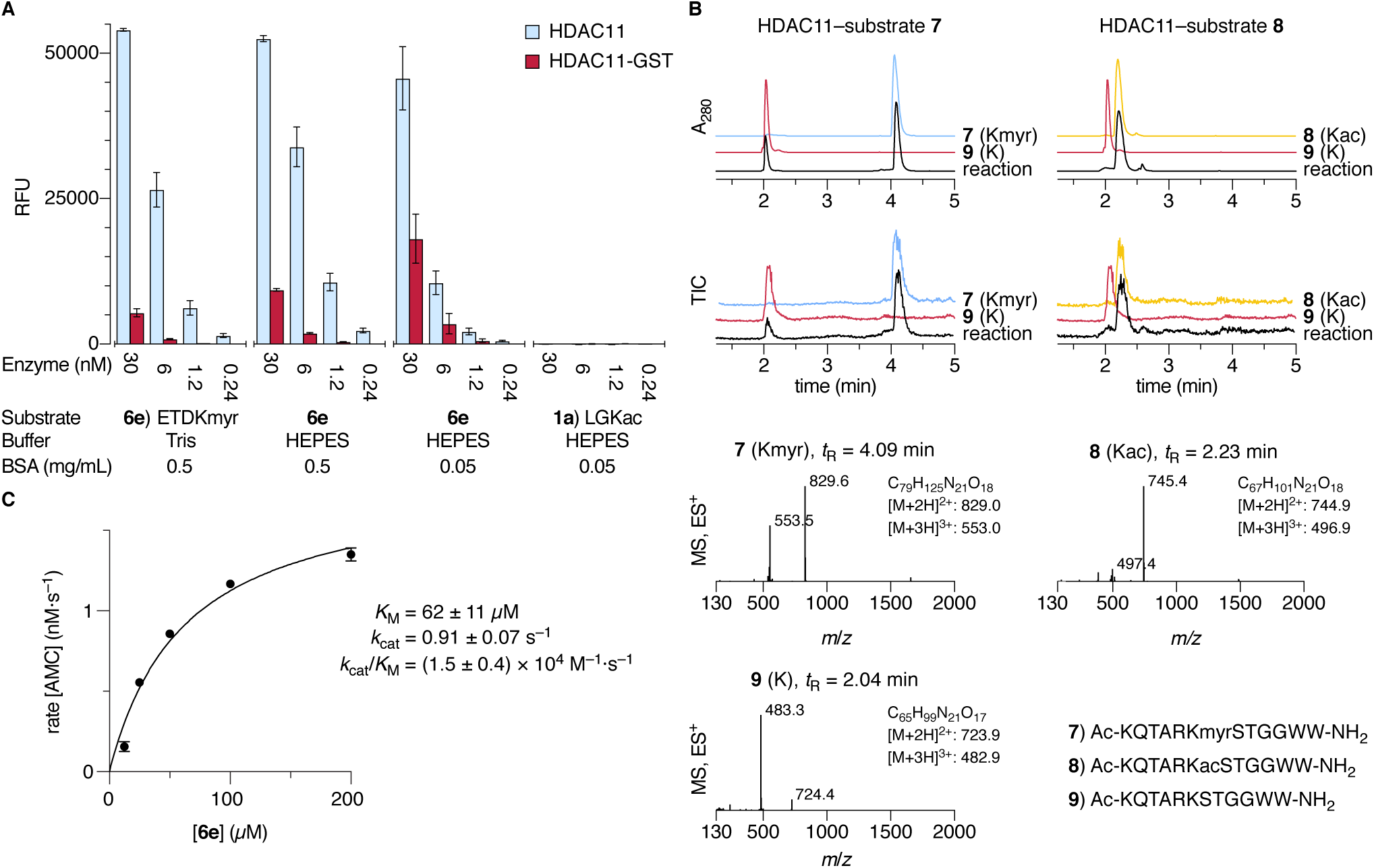
Investigation of HDAC11 demyristoylation efficiency and kinetics. (A) Bar graphs showing the deacylation activity of two HDAC11 constructs at varying enzyme concentrations in different buffers. The results are based on at least two individual end-point assays performed in duplicate. Data are represented as mean ± SD. The enzyme concentrations were based on BCA assays corrected for purity as estimated from Coomassie-stained SDS PAGE gel electrophoresis (Supplemental Fig. S2A) (B) HPLC-MS data of HDAC11-mediated deacylation using non-fluorescent peptide substrates; UV (A_280_) and TIC (ES^+^) chromatograms and mass spectra after 60 min incubation allowing identification of acylated peptide substrates (**7** and **8**) and deacylated peptide product (**9**). (C) Michaelis-Menten plot of steady-state rate experiments for demyristoylation of substrate **6e** at varying concentrations. Data are represented as mean ± SEM.

Full conversion of Kmyr substrate (**6e**) for most enzyme concentrations, which may affect the kinetic profile, renders it difficult to compare the efficiencies based on these end-point data. Thus, we also performed a continuous experiment in the different buffers to determine initial rates (Supplemental Fig. S2B). The HEPES buffer (pH 7.4) containing 0.05 mg/mL of BSA gave rise to the highest initial rates and, because this also has a more relevant pH, we chose this buffer for evaluation of inhibitors and additional substrates. As also indicated in the initial screen, the deacetylation activity of the more active untagged HDAC11 was negligible under these conditions as well (Fig. 2A). Though we cannot rule out that HDAC11 also targets specific Kac-containing substrates in vivo this strongly suggests that HDAC11 is a lysine long chain deacylase enzyme rather than a deacetylase.

Finally, because fluorophore-coupled substrates have been questioned in several contexts (Kaeberlein et al., 2005; Toro et al., 2017), we validated the preference for demyristoylation over deacetylation using non-fluorogenic peptides (**7**–**9**) in an LC-MS-based assay (Fig. 2B). After incubation for 60 minutes with 20 nM of HDAC11, we observed robust conversion of approximately 25% of the myristoylated substrate **7**, while mass spectrometry analysis of the reaction with Kac-containing substrate **8** did not show any trace of product **9** (Fig. 2B).

### Kinetic investigation of HDAC11 demyristoylation

To further characterize this enzyme activity, we investigated the demyristoylation kinetics of untagged HDAC11 by determining initial rates at varying substrate concentrations. We employed continuous assay conditions recently developed for SIRT2 demyristoylation (Galleano et al., 2016), except, to achieve measurable rates at the lower substrate concentrations, we had to remove BSA entirely, an observation that is not surprising given the potential of BSA to bind lipidated peptides. Contrary to the previously characterized demyristoylation activity of SIRT2 (Teng et al., 2015; Galleano et al., 2016), however, the remarkably high enzymatic efficiency for a KDAC measured here (*k*_cat_/*K*_M_ = 1.5 × 10^4^ M^−1^s^−1^) was a result of a high *k*_cat_ rather than a particularly low *K*_M_ (60 μM) (Fig. 2C).

### The HDAC11 demyristoylation activity is inhibited by hydroxamic acid-containing macrocycles but not SAHA (vorinostat)

Previously, assays employing HDAC11 have been shown to require higher enzyme loading (>100 nM) (Madsen and Olsen, 2012a; Madsen et al., 2014; Maolanon et al., 2014; Villadsen et al., 2014; Kitir et al., 2017) using Kac substrates that we now show to be very poor for this enzyme. This leaves open the possibility that previously measured activities may have arisen from minute amounts of co-purified KDACs from the expression organism. We therefore tested a select panel of HDAC inhibitors including approved drugs **SAHA** (vorinostat) (Marks and Breslow, 2007) and **FK-228** (romidepsin) (Furumai et al., 2002) as well as a number of cyclic tetrapeptides (Maolanon et al., 2017) and a trifluoromethylketone previously shown to inhibit all zinc-dependent isozymes (Madsen and Olsen, 2016) (Supplemental Fig. S3). Interestingly, the only compounds that potently inhibited HDAC11 were trapoxin A (**TpxA**) and hydroxamic acid-containing macrocyclic peptides (such as **ApiA^Asuha^**) inspired by archetypal natural product HDAC inhibitors (Fig. 3 and Supplemental Fig. S3) (Furumai et al., 2001; Kitir et al., 2017). Neither romidepsin nor **SAHA** exhibited significant effects at concentrations up to 100 μM, and TSA exhibited considerably lower potency than against class I HDACs (Fig. 3 and Supplemental Fig. S3). Surprisingly, the hydroxamic acid-containing inhibitor **ApiA^Asuha^** rather than epoxyketone **TxpA** exhibited slow, tight-binding kinetics, rendering HDAC11 more similar to HDAC6 than the class I isozymes in this respect (Kitir et al., 2017). On the other hand, macrocycles are generally more potent inhibitors of class I than class IIb isozymes. Although counter-intuitive that the epoxide electrophile-containing inhibitor should exhibit fast-on–fast-off kinetics, this behavior has been supported by co-crystal structures of both HDAC6 and HDAC8 with inhibitors showing intact epoxide moieties (Hai and Christianson, 2016; Porter and Christianson, 2017).

**Figure 3.**
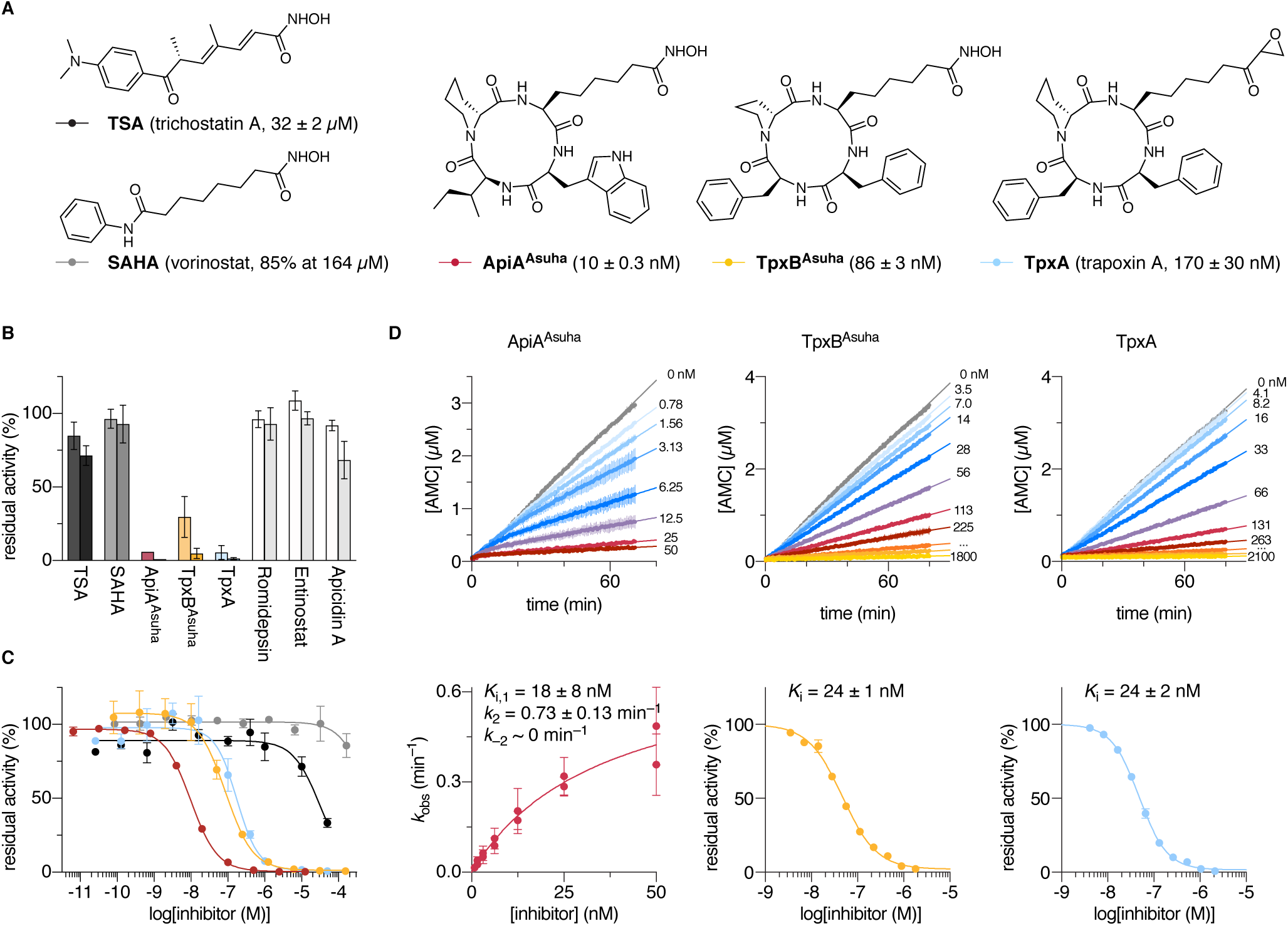
Structures of selected HDAC inhibitors and dose–response curves for inhibition of HDAC11. (A) Chemical structures of trichostatin A (TSA) and vorinostat (SAHA), hydroxamic acid-containing macrocycles (ApiA^Asuha^ and TpxB^Asuha^) and trapoxin A. (B) Bar graphs showing the inhibitory effect on HDAC11 activity of HDAC inhibitors measured at 1 µM and 10 µM inhibitor concentrations. Data are represented as mean ± SD. See also Fig. S3. (C) Concentration–response curves of HDAC11 inhibition by the inhibitors shown in panel A, the obtained IC_50_-values are shown in panel A. Data are represented as mean ± SD. (D) Progression curves and data fitting for HDAC11 inhibition by macrocyclic inhibitors shown in panel A. Data are represented as mean ± SEM. The results are based on at least two individual end-point assays performed in duplicate.

Being able to inhibit this newly discovered enzyme activity is encouraging as it provides further evidence that the effect is a bona fide enzyme mediated catalytic event. Furthermore, the macrocyclic scaffolds may be modularly modified and thus provides an excellent starting point for further optimization of inhibitor potency and selectivity. Combined with the novel assay conditions developed herein, these collective findings provide impetus for further screening regimes aimed at identifying novel chemotypes able to inhibit HDAC11.

### HDAC11 cleaves a narrow selection of long chain acyl groups

Finally, we addressed the substrate scope with respect to the identity of the acyl modification by incorporating various lengths of fatty acids as well as a selection of oxidized and unsaturated versions (Fig. 4A). This screen showed that HDAC11 preferably cleaves fatty acid-based modifications with 12– 14 carbon atoms in length, while both truncation (**4u**, **5c**, **5d**, **5u**, **5v**, and **6d**) and elongation (**5y**) give rise to diminished activity. Furthermore, oxidation at the 3-position to give hydroxyl (**6æ**) or ketone (**6ø**) modifications abolished activity while the structurally more similar α, β-unsaturated substrate (**6z**) was cleaved (Fig. 4). This indicates a quite narrow substrate specificity in contrast to another long chain deacylase, SIRT2, which cleaves these modifications with similar activity as Kmyr (Galleano et al., 2016).

**Figure 4.**
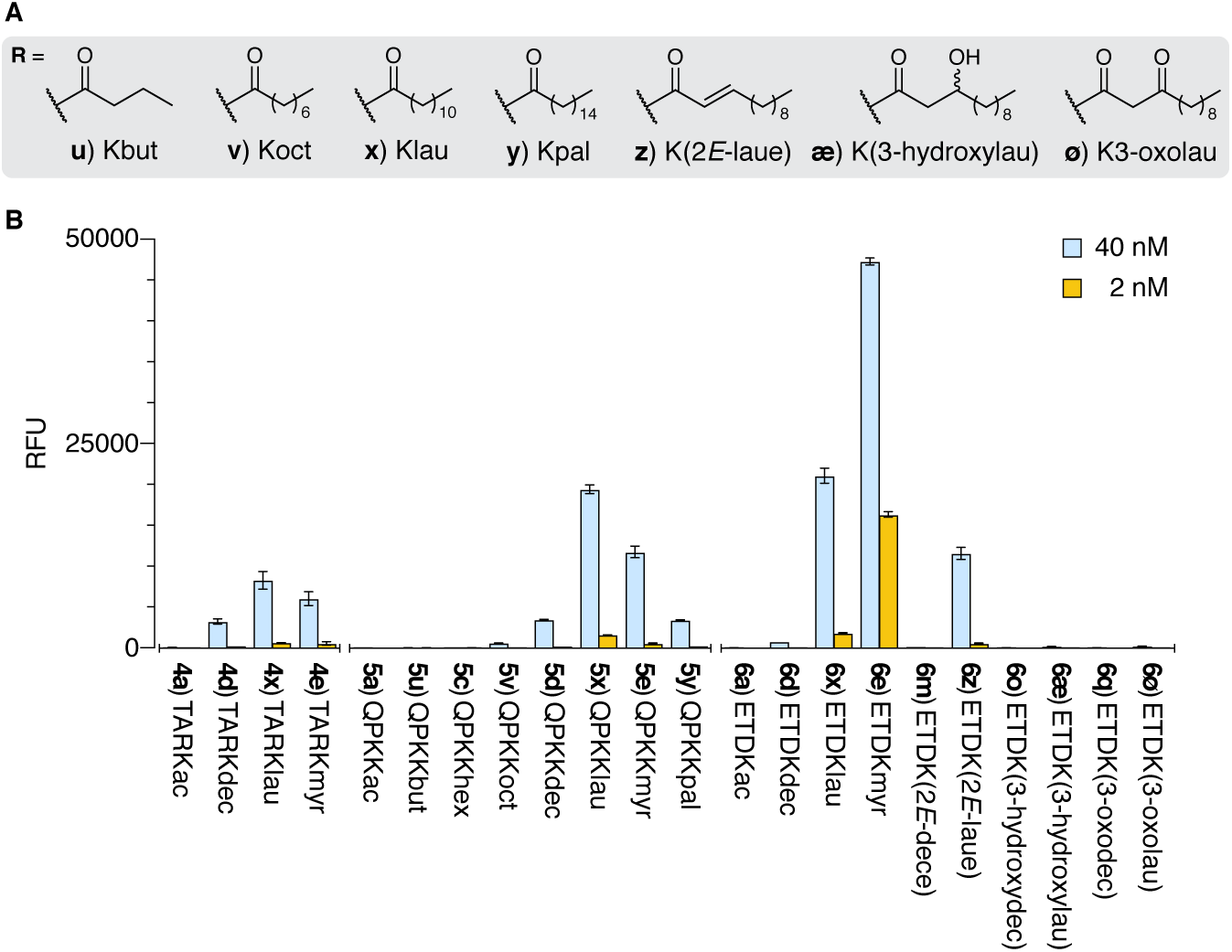
Screening for HDAC11 activity against additional long chain acyl modifications on lysine. (A) Chemical structures of the additional side chains included in the substrate series. (B) Bar graphs showing the conversion of substrates in end-point assays (given as relative fluorescence units). Initial concentrations of substrates were 50 μM and the enzyme loading was either 2 nM or 40 nM of recombinant, full-length, untagged HDAC11. The results are based on at least two individual end-point assays performed in duplicate. Data are represented as mean ± SD.

Interestingly, we observed a distinct effect on conversion depending on the peptide sequence, where the negatively charged ETDK-based peptide (scaffold **6**) was turned over to a significantly higher degree than the two positively charged substrates (scaffolds **4** and **5**). Unfortunately, non-charged substrates of this type containing the Kmyr modification are not soluble in aqueous media, and cannot be commented on in this context. However, more thorough investigation of the importance of peptide sequence will be an interesting subject to explore in the future.

## DISCUSSION

We have performed an extensive screening for potential enzymatic activity of the human zinc-dependent HDACs against a wide range of ε-*N*-acyllysine modifications. Mostly, this did not reveal any surprises, *i.e.*, class I HDACs as well as HDAC6 robustly removed acetyl groups and HDAC1–3 were able to cleave Kcr as well as other short chain modifications with structural similarity. Class IIa enzymes (Lahm et al., 2007) as well as HDAC3 (Madsen and Olsen, 2012a), HDAC8 (Bradner et al., 2010), and HDAC11 (Inks et al., 2012) removed the physiologically non-relevant trifluoroacetamide (Ktfa). However, the one result that stood out was the efficient cleavage of substrate **6e** (Ac-ETDKmyr-AMC) by HDAC11, because long chain deacylation by this isozyme is unprecedented. Histone deacetylase 11 has been implicated in immune response cascades and shown to regulate the expression of the anti-inflammatory cytokine interleukin 10 (IL-10) by association with HDAC6 (Villagra et al., 2009). More recently, HDAC11 has also been shown to be involved on Fox3p+ T-regulatory cell function (Huang et al., 2017), but it remains the least understood isozyme of the class. Demonstration of this novel substrate selectivity is therefore of major importance for the continued investigation of HDAC11 function. Thus, we examined this activity in more detail, first by comparing different recombinant enzyme constructs, showing that the untagged full-length protein was significantly more active than its GST-tagged fusion protein. We also tested different buffers to establish ideal conditions for in vitro evaluation of HDAC11. Then, we validated the remarkable selectivity for Kmyr over Kac using dodecamer peptides in an LC-MS assay to demonstrate that the novel activity was not an artifact resulting from the AMC fluorophore, and further showed selectivity for long chain deacylation using an extended series of AMC-conjugated peptides.

We determined kinetic parameters, which furnished remarkable enzymatic efficiency (*k*_cat_/*K*_M_ = 1.5 × 10^4^ M^−1^s^−1^) for an HDAC or sirtuin enzyme using fluorescence-based assays. This high efficiency combined with the narrow substrate scope strongly indicates that lysine demyristoylation is a bona fide enzymatic activity of HDAC11. However, we cannot exclude the possibility that HDAC11 may also be able to hydrolyze Kac under certain conditions such as in a chromatin complex for example. It has been shown to associate with HDAC6 (Gao et al., 2002; Cheng et al., 2014) and, since one is primarily cytosolic (HDAC6) whereas the other is primarily nuclear (HDAC11), it may be speculated that both can shuttle between the two compartments and that their activities vary depending on environment.

In addition to raising new questions, the present work importantly provides reliable protocols for testing inhibitors of HDAC11 for the first time and, surprisingly, most archetypical HDAC inhibitors proved inefficient. Among the compounds tested in the present study, only cyclic tetrapeptides with strong zinc-binding groups potently inhibited HDAC11 demyristoylase activity. In agreement with the original account describing this enzyme, trapoxin A (**TpxA**) was a potent inhibitor (Gao et al., 2002), and so were hydroxamic acid-containing versions of select macrocycles (**TpxB^Asuha^** and **ApiA^Asuha^**). Although these inhibitors are not HDAC11-selective as they have all been shown to potently inhibit several other members of the enzyme class (Kitir et al., 2017), it was encouraging to identify potent inhibitors that may undergo further optimization. The significant difference in affinity depending on macrocyclic scaffold recorded for the two hydroxamates, combined with the substrate selectivity between peptide scaffolds, provide hope that structure–activity relationship studies may lead to improved selectivity. The robust and enzyme-efficient assay protocols developed in this work, employing either fluorescence or LC-MS, will be instrumental in the accurate evaluation of HDAC11 inhibitors in the future.

## SIGNIFICANCE

Histone deacetylase 11 is the least characterized of the human zinc-dependent HDACs. Here we provide novel insight into its enzymology at the molecular level by addressing its substrate specificity, enzyme kinetics, and susceptibility to a range of standard HDAC inhibitors in vitro. Based on our data, HDAC11 is the first isozyme of this class demonstrated to exhibit a preference for physiologically relevant acyl groups other than acetyl. Moreover, the identified activity against lysine side chains modified with the 14-carbon atom myristic acid, which is a known PTM, was remarkably efficient for an HDAC enzyme. This new substrate specificity for HDAC11 will have major implications for investigation of its biological function and has already been shown to be of relevance in a cellular context (personal communication with Prof. H. Lin). In addition, we developed enzyme-economical and reliable assay protocols that allow for the characterization of potential inhibitors of HDAC11. Surprisingly, a number of HDAC inhibitors previously thought to inhibit this enzyme proved ineffective, and the few compounds that inhibited HDAC11, exhibited interesting inhibition kinetics, which will be instructive in the future search for more potent and selective tools compounds to help study HDAC11. This work will be key to unlocking the elusive function of HDAC11 in immune response and anti-inflammatory cascades and, potentially, in additional biological mechanisms.

### STAR*METHODS

Detailed methods are provided in the online version of this paper and include the following:

- KEY RESOURCES TABLE
- CONTACT FOR REAGENT AND RESOURCE SHARING
- METHOD DETAILS
  - Buffers used for in vitro enzymatic characterization
  - Substrate screening assays
  - End-point inhibition assays
  - Continuous assays
  - Michaelis-Menten assays
  - HPLC-MS-based assays
  - Chemical synthesis

## AUTHOR CONTRIBUTIONS

Conceptualization, C.A.O.; Investigation, C.M.-Y., I.G. and A.S.M.; Writing – Original Draft, C.A.O.; Writing – Review & Editing, C.M.-Y., I.G., A.S.M., and C.A.O.; Visualization, A.S.M.; Supervision, A.S.M. and C.A.O.; Funding Acquisition, C.A.O.; Project Administration, C.A.O.

## ACKNOWLEDGMENTS

We thank Dr. A. M. Maolanon and Dr. R. G. Ohm for synthesis of macrocyclic inhibitors. We gratefully acknowledge financial support from the University of Copenhagen (partial PhD fellowships for I.G. and C.M.-Y.), The Carlsberg Foundation (2011_01_0169, 2013_01_0333, and CF15-0115; CAO), The Novo Nordisk Foundation (NNF15OC0017334; CAO), and the European Research Council (ERC-CoG-725172 – *SIRFUNCT*; C.A.O.).

## Table of Content Graphics

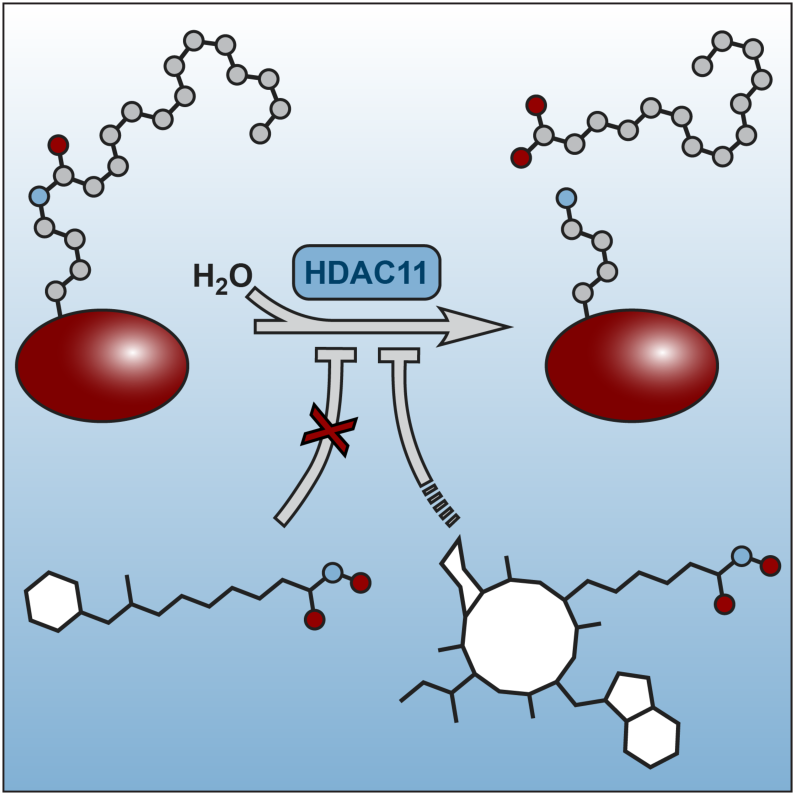

## STAR*METHODS

### CONTACT FOR REAGENT AND RESOURCE SHARING

Further information and requests for resources and reagents should be directed to and will be fulfilled by the Lead Contact, Christian Adam Olsen.

## METHOD DETAILS

### Buffers used for in vitro enzymatic characterization

- Tris buffer was prepared as described in Biomol International product sheets BML-KI-143 [http://www.enzolifesciences.com/BML-AK500/fluorde-lys-hdac-fluorometric-activity-assay-kit/] [50 mM Tris/Cl, 137 mM NaCl, 2.7 mM KCl, MgCl_2_ 1 mM, 0.5 mg/mL BSA [A7030], pH 8.0].
- HEPES buffer was prepared as previously described(Bradner et al., 2010; Galleano et al., 2016) [50 mM HEPES/Na, 100 mM KCl, 0.001% (v/v) Tween-20, 0.05 mg/mL BSA, 200 μM tris(2-carboxyethyl)phosphine (TCEP), pH 7.4]. The amount of BSA added is specified for each type of assay.

All fluorescence-based HDAC deacylase activity assays were performed in black low binding 96-well microtiter plates (Corning half-area wells), with duplicate series in each assay and each assay performed at least twice. Control wells without enzyme were included in each plate. All reactions were performed in assay buffer (see above) with appropriate concentrations of substrates and inhibitors obtained by dilution from 2–200 mM stock solutions in milliQ water or DMSO, and appropriate concentration of enzyme obtained by dilution of the stock provided by the supplier. Data analysis was performed using GraphPad Prism 7.0.

#### Substrate screening assays

The initial screening for substrate deacylation activity was performed in Tris buffer with end-point fluorophore release by trypsin. For a final volume of 25 μL per well, acyl substrates (50 μM) were added to each well followed by a solution of the appropriate HDAC enzyme (50 nM, GST tagged HDAC11 at 73 nM was used for HDAC11 data). The reaction was incubated at 37 °C for 60 min, then a solution of trypsin (25 μL, 5.0 mg/mL; final concentration of 2.5 mg/mL) was added, and the assay development was allowed to proceed for 90 min at room temperature before fluorescence analysis. The data were analyzed to afford [AMC] relative to control wells.

HDAC11 titration in different buffers (Tris or HEPES, with 0.05 or 0.5 mg/mL BSA [A7030], Figure 2A) was performed in a similar manner, with incubation of acyl substrate (ETDKmyr or LGKac, 50 µM) and HDAC11 (untagged or GST tagged, 0.24–30 nM) at 37 °C for 60 min, and development with a solution of trypsin (25 µL, 0.4 mg/mL; final concentration of 0.2 mg/mL).

The screening for long chain substrate deacylation activity was performed in the same manner in HEPES buffer (0.05 mg/mL BSA [A7030]) and HDAC11 enzyme (untagged, 2 nM and 40 nM concentrations).

### End-point inhibition assays

End-point inhibition assays (concentration–response) were performed in HEPES buffer (with 0.05 mg/mL BSA [A7030]) in a final volume of 25 μL per well, where inhibitor (5-fold dilution series) was incubated with Ac-ETDKmyr (**6e**, 50 µM) and HDAC11 (untagged, 1 nM) for 30 min at 37 °C. Thereafter, a solution of trypsin (25 µL, 0.4 mg/mL; final concentration of 0.2 mg/mL) was added, and the assay development was allowed to proceed for 15 min at room temperature before fluorescence analysis. The data were analyzed to afford residual activity relative to control wells, and assuming a standard fast-on/fast-off mechanism, IC_50_ values were obtained by fitting the resulting data to the concentration–response equation with variable Hill slope (**Eq. 1**).

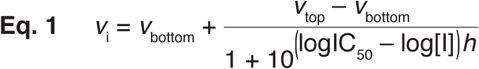

### Continuous assays

Continuous inhibition assays (concentration-response) were performed in HEPES buffer (with 0.05 mg/mL BSA [A3059]) in a final volume of 50 µL per well, where dilution series of the inhibitor (2-fold dilutions) were incubated with Ac-ETDKmyr (**6e**, 60 µM), trypsin (100 ng/µL) and HDAC11 (untagged, 0.5 nM). In situ fluorophore release was monitored immediately by fluorescence readings recorded continuously every 30 s for 100 min at 25 °C. The data were fitted to the relevant equations (**Eq. 2** or **3**) to obtain either initial linear rates (*ν*) or apparent first-order rate constant (*k*_obs_) for each inhibitor concentration. For fast-on–fast-off inhibitors (**TpxB^Asuha^** and **TpxA**), secondary plots were then fitted to **Eq. 1** and the obtained IC_50_ values converted to *K*_i_ values by the Cheng-Prusoff equation (**Eq. 4**). For slow binding inhibitors (**ApiA^Asuha^**), secondary plots were fitted to the relevant equation (**Eq. 5**) to obtain dissociation constant (*K*_i,1_) and kinetic parameters (*k*_2_ and *k*_–2_). For further discussion of the applied equations, please see (Kitir et al., 2017).

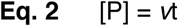

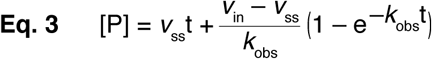

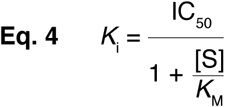

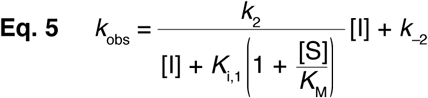

Continuous assays (Figure S2) were performed in different buffers (BSA [A3059]) in a final volume of 50 µL per well, where acyl substrate (20 µM) was incubated with HDAC11 (untagged, 2 nM) and different concentrations of trypsin (200–25 ng/µL). In situ fluorophore release was monitored immediately by fluorescence readings recorded continuously every 30 s for 100 min at 25 °C.

#### Michaelis-Menten assays

Rate experiments for determination of kinetic parameters were performed in HEPES buffer (no BSA added) in a final volume of 50 μL per well, where Ac-ETDKmyr (**6e**, 2-fold dilutions) was incubated with trypsin (200 ng/μL) and HDAC11 (untagged, 2 nM). In situ fluorophore release was monitored immediately by fluorescence readings recorded continuously every 30 s for 100 min at 25 °C to obtain initial rates ν_0_ (nM⋅s^−1^) for each concentration. The data were fitted to the Michaelis−Menten equation to afford *K*_*m*_ (μM) and *k*_*cat*_ (s^−1^) values.

#### HPLC-MS-based assays

After incubating HDAC11 (untagged, 20 nM) and the relevant dodecapeptide substrate (**7** [Kac] or **8** [Kmyr], 50 µM) for 60 min at 37 °C, a sample of the reaction mixture (25 µL) was taken out and quenched by addition of MeOH/HCOOH (94:6 (v/v), 12.5 µL). The samples were analyzed by HPLC-MS on a Waters Acquity ultra-HPLC-MS system equipped with a diode array detector. A gradient with eluent I (0.1% HCOOH in water (v/v)) and eluent II (0.1% HCOOH in acetonitrile (v/v)) rising linearly 5–100% during t = 0.10–8.00 min was applied at a flow rate of 0.6 mL/min. The obtained chromatograms at 280 nm were used to determine reaction progression, and the obtained mass spectra used to determine formation of the desired deacylated dodecamer product (**9** [K]).

#### Protein concentration assay

##### HDAC1-10 Stock solution concentrations were used as provided by the vendor

HDAC11 protein concentration was determined with the bicinchoninic acid (BCA) protein concentration assay. Protein solutions (20 µL) were mixed with BCA reagent mixture (160 µL), incubated 30 min at 37 °C and read for absorption at 562 nm. The standard curve (0.016 to 1.00 µg/µL) was prepared with BSA protein standards in triplicate. HDAC11 commercial Stock solutions were diluted 1:10 (untagged) or 1:20 (GST-tag) in MilliQ water, analyzed in triplicate, and total protein concentration was calculated with respect to the standard curve. HDAC11 relative purity was determined by coomasie-stained gel electrophoresis (Supplemental Fig. S2A) and used for the calculation of HDAC11 protein concentration in each Stock solution. HDAC11 untagged: 0.79 mg/mL, HDAC11 GST-tag: 2.43 mg/mL.

#### Chemical synthesis

All reagents and solvents were of analytical grade and used without further purification as obtained from commercial suppliers. Anhydrous solvents were purchased or prepared according to literature.(Williams and Lawton, 2010) Reactions were conducted under an atmosphere of nitrogen whenever anhydrous solvents were used. All reactions were monitored by thin-layer chromatography (TLC) using silica gel coated plates (analytical SiO_2_-60, F-254). TLC plates were visualized under UV light and by dipping in either (a) a solution of potassium permanganate (10 g/L), potassium carbonate (67 g/L) and sodium hydroxide (0.83 g/L) in water, (b) a solution of ninhydrin (3 g/L) in 3% acetic acid in water (v/v), or (c) a solution of vanillin (5 g/L) in 80% sulfuric acid in ethanol (v/v) followed by heating with a heat gun. Evaporation of solvents was carried out under reduced pressure at temperatures below 45 °C. ^1^H NMR and ^13^C NMR were recorded on a Bruker Avance III HD equipped with a cryogenically cooled probe, at 600 MHz and 151 MHz, respectively. Chemical shifts are reported in ppm relative to deuterated solvent as internal standard (δ_H_: DMSO-d_6_ 2.50 ppm; δ_C_: DMSO-d_6_ 39.52 ppm). Coupling constants are reported in Hz. Assignment of NMR spectra are based on correlation spectroscopy (COSY, HSQC, and HMBC spectra). Loading of resin during solid phase peptide synthesis was checked spectrophotometrically, quantifying the amount of Fmoc released upon cleavage of a small sample.(Gude et al., 2002) UPLC-MS analyses were performed on a Waters Acquity ultra high-performance liquid chromatography system equipped with a C18 Phenomenex Kinetex column [50 mm × 2.1 mm, 1.7 μm, 100 Å], using a gradient of eluent I (0.1% HCOOH in water) and eluent II (0.1% HCOOH in acetonitrile) rising linearly from 0% to 95% of eluent II during t = 0.00−5.20 min. Analytical reversed-phase HPLC was performed on an Agilent 1100 LC system equipped with a C18 Phenomenex Kinetex column [150 mm × 4.60 mm, 2.6 μm, 100 Å] and a diode array UV detector, using a gradient of eluent III (water−MeCN−TFA, 95:5:0.1) and eluent IV (0.1% TFA in MeCN) rising linearly eluent IV as stated for each compound, with a flow rate of 1 mL/min.

#### Ac-Leu-Gly-Lys(acyl)-AMC substrates

See (Madsen and Olsen, 2012b). Briefly, Ac-Leu-OH was coupled to H-Gly-OMe using HOBt. The dipeptide was then coupled to H-Lys(Boc)-AMC using HOBt/DIC, upon hydrolysis of the methyl ester. Removal of the Boc protection-group of the resulting tripeptide in presence of CF_3_COOH afforded Ac-Leu-Gly-Lys-(7-amino-4-methyl-coumarin) (**S1**) as the corresponding trifluoroacetate salt, which was then acylated and purified by preparative HPLC to afford the desired products.

#### Ac-Gln-Pro-Lys-Lys(acyl)-AMC substrates

See (Madsen et al., 2016). Briefly, Fmoc-Lys-OAlloc was loaded on 2-chlorotrityl linker polystyrene resin. The tripeptide Ac-Gln(Trt)-Pro-Lys(resin)-OAlloc was assembled on resin using standard solid phase peptide synthesis (SPPS). The tripeptide on resin was coupled to the fluorogenic H-Lys(Teoc)-AMC using DIC, upon Alloc-deprotection. The tetrapeptide was Teoc deprotected on resin, acylated and eventually deprotected and cleaved from resin at the same time in presence of CF_3_COOH. Preparative HPLC purification afforded the desired products.

#### Ac-Glu-Thr-Asp-Lys(acyl)-AMC substrates

See (Galleano et al., 2016). Briefly, Ac-Glu(^*t*^Bu)-Thr(^*t*^Bu)-Asp(^*t*^Bu)-OH was assembled on 2-chlorotrityl linker polystyrene resin via standard SPPS. The tripeptide was cleaved from the resin under mild conditions and coupled to TFA⋅H-Lys(Fmoc)-AMC, followed by Fmoc deprotection, affording the appropriately protected tetrapeptide Ac-Glu(^*t*^Bu)-Thr(^*t*^Bu)-Asp(^*t*^Bu)-Lys-AMC (**S2**). The tetrapeptide was acylated according to literature, upon. Deprotection in presence of CF_3_COOH and preparative HPLC afforded the desired compounds.

#### General acylation procedure for the compound 6 series

The desired acid (0.06 mmol) was dissolved in anhydrous CH_2_Cl_2_ (2 mL). Then HATU (23 mg, 0.06 mmol) and lutidine (12 mg, 0.12 mmol) were added and the resulting suspension was stirred for 10 min, before the mixture was cooled to 0 °C and Ac-Glu(^*t*^Bu)-Thr(^*t*^Bu)-Asp(^*t*^Bu)-Lys-AMC(Galleano et al., 2016) (**S2**, 40 mg, 0.05 mmol) was added. After the reaction reached completion as judged by LC-MS analysis, the solution was partitioned between CH_2_Cl_2_ (6 mL) and brine (5 mL). The aqueous phase was back extracted with CH_2_Cl_2_ (3 × 5 mL), and the combined organic layer was washed with aqueous HCl (0.5 M, 2 × 10 mL). The acidic layer was back extracted with CH_2_Cl_2_ (2 × 8 mL), and the combined organic phase was washed with sat aqueous NaHCO_3_ (2 × 10 mL). The basic aqueous phase was also extracted with CH_2_Cl_2_ (2 × 10 mL), and the resulting combined organic phase was washed with brine (30 mL). Again, the aqueous phase was back extracted with CH_2_Cl_2_ (2 × 10 mL), and the final combined organic layer was dried over MgSO_4_, filtered, and concentrated in vacuo to give a residue, which was stirred in CF_3_COOH−CH_2_Cl_2_−H_2_O (2 mL, 50:48:2) for 90 min. The solution was then concentrated under reduced pressure and the crude residue was precipitated from ice-cold diethyl ether (15 mL). The crude residue was purified by reversed-phase preparative HPLC to afford the desired product.

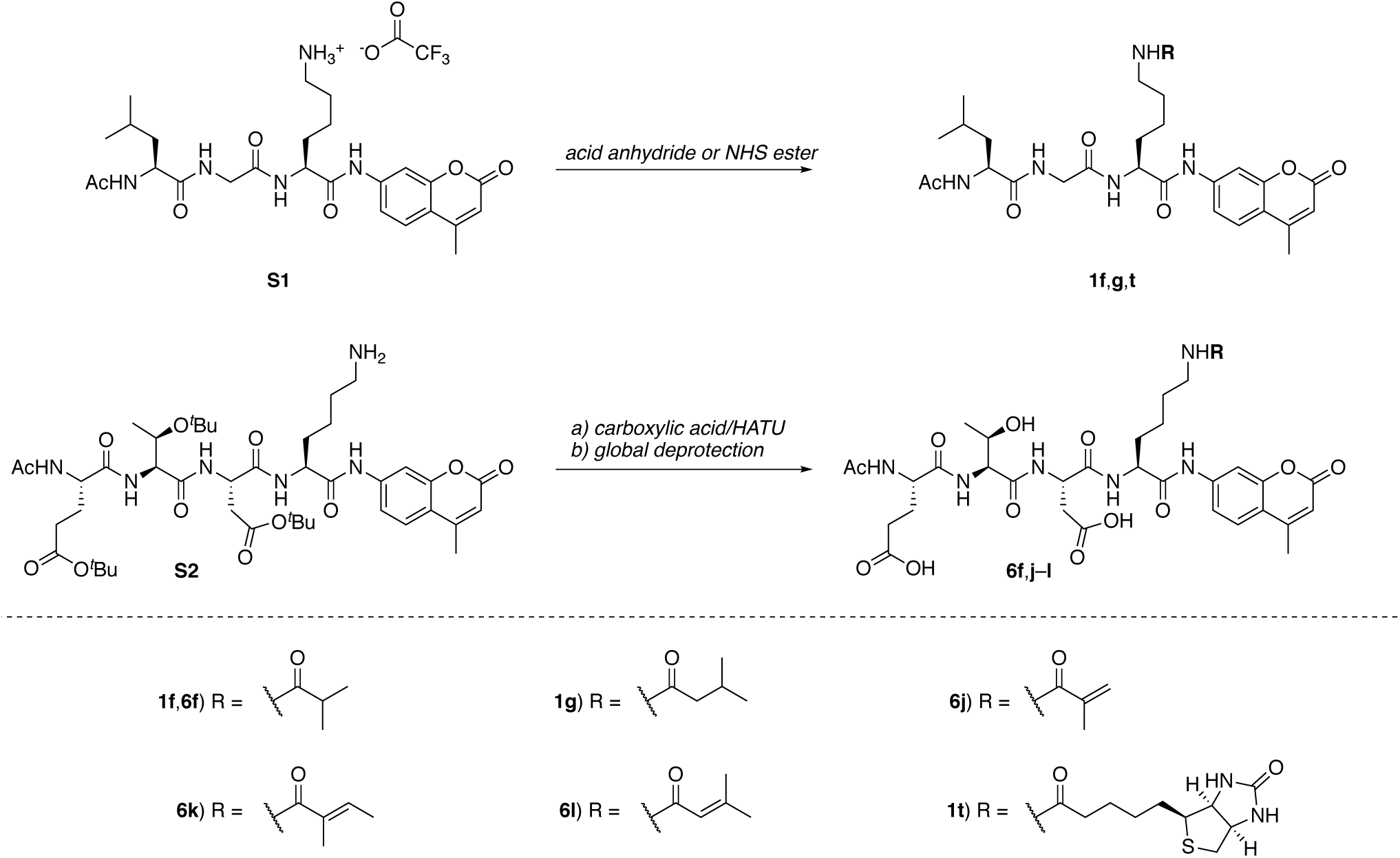

#### Ac-Leu-Gly-Lys(isobutyryl)-AMC

(**1f**). iPr_2_NEt (10.3 µL, 60 µmol) and isobutyric anhydride (5.9 µL, 36 µmol) was dissolved in anh CH_2_Cl_2_ (0.5 mL), followed by addition of Ac-Leu-Gly-Lys-(7-amino-4-methyl-coumarin) trifluoroacetate salt(Madsen and Olsen, 2012b) (15 mg, 24 µmol). After stirring at rt for 40 min, MeOH (0.5 mL) was added, and the reaction mixture evaporated to dryness. The resulting residue was purified by preparative HPLC, affording Ac-Leu-Gly-Lys(isobutanoyl)-(7-amino-4-methyl-coumarin) (9.0 mg, 65%) as a white fluffy material. ^1^H NMR (600 MHz, DMSO-*d*_6_) δ 10.36 (s, 1H, NH_AMC_), 8.30 (t, *J* = 5.9 Hz, 1H, NH_gly_), 8.08 (d, *J* = 7.3 Hz, 1H, NH_leu_), 8.01 (d, *J* = 7.6 Hz, 1H, NHα_lys_), 7.79 (d, *J* = 2.0 Hz, 1H, H8_AMC_), 7.69–7.74 (m, 1H, H5_AMC_), 7.66 (t, *J* = 5.7 Hz, 1H, NHε_lys_), 7.52 (dd, *J* = 8.8, 2.0 Hz, 1H, H6_AMC_), 6.24–6.28 (m, 1H, H3_AMC_), 4.34–4.40 (m, 1H, Hα_lys_), 4.19–4.25 (m, 1H, Hα_leu_, overlap with solvent signal), 3.68–3.77 (m, 2H, Hα_gly_, overlap with solvent signal), 2.98–3.03 (m, 2H, Hε_lys_, overlap with solvent signal), 2.40 (d, *J* = 1.5 Hz, 3H, 4_AMC_-CH_3_), 2.30 (hept, *J* = 6.8 Hz, 1H, NHε_lys_COC*H*), 1.85 (s, 3H, C*H*_*3*_CONH_leu_), 1.69–1.78 (m, 1H, Hβ_lys,A_), 1.55–1.69 (m, 2H, Hγ_leu_,Hβ_lys,B_), 1.42–1.50 (m, 2H, Hβ_leu_), 1.36–1.42 (m, 2H, Hδ_lys_), 1.21–1.36 (m, 2H, Hγ_lys_), 0.954 (d, *J* = 6.8 Hz, 3H, NHε_lys_COCH(C*H*_*3*_)_2,A_), 0.949 (d, *J* = 6.9 Hz, 3H, NHε_lys_COCH(C*H*_*3*_)_2,B_), 0.88 (d, *J* = 6.6 Hz, 3H, CH_3,leu,A_), 0.83 (d, *J* = 6.6 Hz, 3H, CH_3,leu,B_). ^13^C NMR (151 MHz, DMSO) δ 175.9 (CONHε_lys_), 172.9 (CO_leu_), 171.4 (CO_lys_), 169.7 (CO_ac_), 169.0 (CO_gly_), 160.0 (C2_AMC_), 153.6 (C8a_AMC_), 153.1 (C4_AMC_), 142.1 (C7_AMC_), 125.9 (C5_AMC_), 115.3 (C6_AMC_), 115.1 (C4a_AMC_), 112.3 (C3_AMC_), 105.8 (C8_AMC_), 53.6 (Cα_lys_), 51.5 (Cα_leu_), 42.0 (Cα,_ly_), 40.5 (Cβ_leu_), 38.1 (Cε_lys_), 34.0 (NHε_lys_CO*C*H(CH_3_)_2_), 31.4 (Cβ_lys_), 28.8 (Cδ_lys_), 24.2 (Cγ_leu_), 22.9 (CH_3,leu,A_), 22.8 (Cγ_lys_), 22.5 (*C*H_3_CO), 21.6 (CH_3,leu,B_), 19.6 (NHε_lys_COCH(*C*H_3_)_2_), 18.0 (4_AMC_-CH_3_). HRMS *m*/*z* calcd for C_30_H_44_N_5_O_7_^+^ [M+H]^+^, 586.3235; found, 586.3227.

#### Ac-Leu-Gly-Lys(isovaleryl)-AMC

(**1g**). iPr_2_NEt (10.3 µL, 60 µmol) and isovaleric anhydride (3.4 µL, 29 µmol) was dissolved in anh CH_2_Cl_2_ (0.5 mL), followed by addition of Ac-Leu-Gly-Lys-(7-amino-4-methyl-coumarin) trifluoroacetate salt(Madsen and Olsen, 2012b) (15 mg, 24 µmol). After stirring at rt for 40 min, MeOH (0.5 mL) was added, and the reaction mixture evaporated to dryness. The resulting residue was purified by preparative HPLC, affording Ac-Leu-Gly-Lys(isovaleryl)-(7-amino-4-methyl-coumarin) (**1g**, 8.4 mg, 60%) as a white fluffy material. ^1^H NMR (600 MHz, DMSO-*d*_6_) δ 10.36 (s, 1H, NH_AMC_), 8.31 (t, *J* = 5.9 Hz, 1H, NH_gly_), 8.08 (dd, *J* = 7.3 Hz, 1H, NH_leu_), 8.00 (d, *J* = 7.5 Hz, 1H, NHα_lys_), 7.79 (d, *J* = 2.0 Hz, 1H, H8_AMC_), 7.69–7.74 (m, 2H, H5_AMC_, NHε_lys_), 7.52 (dd, *J* = 8.7, 2.0 Hz, 1H, H6_AMC_), 6.26 (d, *J* = 3.2 Hz, 1H, H3_AMC_), 4.35–4.40 (m, 1H, Hα_lys_), 4.19–4.24 (m, 1H, Hα_leu_, overlap with solvent signal), 3.68–3.77 (m, 2H, Hα_gly_, overlap with solvent signal), 3.01 (q, *J* = 6.7 Hz, 2H, H_ε,lys_, overlap with solvent signal), 2.40 (d, *J* = 1.9 Hz, 3H, 4_AMC_-CH_3_), 1.87–1.96 (m, 3H, NH_ε,lys_COC*H*_*2*_C*H*), 1.85 (s, 3H, C*H*_*3*_CONH_leu_), 1.70–1.77 (m, 1H, Hβ_lys,A_), 1.57–1.69 (m, 2H, Hγ_leu_,Hβ_lys,B_), 1.42–1.50 (m, 2H, Hβ_leu_), 1.37–1.42 (m, 2H, Hδ_lys_), 1.22–1.37 (m, 2H, Hγ_lys_), 0.88 (d, *J* = 6.6 Hz, 3H, CH_3,leu,A_), 0.84 (d, *J* = 6.6 Hz, 3H, CH_3,leu,B_), 0.819 (d, *J* = 6.3 Hz, 3H NHε_lys_COCH_2_CH(C*H*_*3*_)_2,A_), 0.817 (d, *J* = 6.3 Hz, 3H, NHε_lys_COCH_2_CH(C*H*_*3*_)_2,B_). ^13^C NMR (151 MHz, DMSO) δ 172.9 (CO_leu_), 171.4 (CO_lys_), 171.3 (CONHε_lys_), 169.7 (CO_ac_), 169.0 (CO_gly_), 160.0 (C2_AMC_), 153.6 (C8a_AMC_), 153.1 (C4_AMC_), 142.1 (C7_AMC_), 125.9 (C5_AMC_), 115.3 (C6_AMC_), 115.1 (C4a_AMC_), 112.3 (C3_AMC_), 105.8 (C8_AMC_), 53.6 (Cα_lys_), 51.5 (Cα_leu_), 44.8 (NHε_lys_CO*C*H_2_), 42.0 (Cα_gly_), 40.5 (Cβ_leu_), 38.1 (Cε_lys_), 31.4 (Cβ_lys_), 28.9 (Cδ_lys_), 25.5 (NHε_lys_COCH_2_*C*H), 24.2 (Cγ_leu_), 22.9 (CH_3,leu,A_), 22.8 (Cγ_lys_), 22.5 (*C*H_3_CO), 22.3 (NHε_lys_COCH_2_CH(*C*H_3_)_2_), 21.6 (CH_3,leu,B_), 18.0 (4_AMC_-CH_3_). HRMS *m*/*z* ([M+H]^+^, Calcd). HRMS *m*/*z* calcd for C_31_H_46_N_5_O_7_^+^ [M+H]^+^, 600.3392; found, 600.3383.

#### Ac-Leu-Gly-Lys(biotinyl)-AMC (**1t**)

Ac-Leu-Gly-Lys-(7-amino-4-methyl-coumarin) trifluoroacetate salt(Madsen and Olsen, 2012b) (33 mg, 48 µmol), *N*-succinimidyl biotinate (24 mg, 71 µmol), and iPr_2_NEt (20 µL, 115 µmol) was stirred in anh CH_2_Cl_2_ (1 mL) for 45 min, then MeOH (1 mL) was added and the reaction mixture was evaporated to dryness, then purified by by preparative HPLC to afford desired Ac-Leu-Gly-Lys(biotinyl)-AMC (**1t**, 29 mg, 76%) as a white fluffy material. ^1^H NMR (DMSO-*d*_6_) *δ* 10.36 (s, 1H, NH_AMC_), 8.32 (t, *J* = 5.8 Hz, 1H, NH_gly_), 8.10 (d, *J* = 7.3 Hz, 1H, NH_leu_), 8.01 (d, *J* = 7.6 Hz, 1H, NHα_lys_), 7.79 (d, *J* = 2.0 Hz, 1H, H8_AMC_), 7.74 (t, *J* = 5.7 Hz, 1H, NHε_lys_), 7.72–7.70 (m, 1H, H5_AMC_), 7.53 (dd, *J* = 8.7, 2.0 Hz, 1H, H6_AMC_), 6.43 (br s, 2H, H–N1_biotin_,H–N3_biotin_), 6.26 (d, *J* = 1.8 Hz, 1H, H3_AMC_), 4.37 (m, 1H, Hα_lys_), 4.30 (m, 1H, H6a_biotin_), 4.22 (m, 1H, Hα_leu_), 4.12 (dd, *J* = 7.7, 4.4 Hz, 1H, H3a_biotin_), 3.75 (m_ABX,A_, *J* =16.8, 6.0, 1H, Hα_gly,A_), 3.72 (m_ABX,B_, *J* = 16.8, 6.1, 1H, Hα_gly,B_), 3.07 (ddd, *J* = 8.5, 6.2, 4.4 Hz, 1H, H4_biotin_), 3.01 (q, *J* = 6.6 Hz, 2H, Hε_lys_), 2.81 (dd, *J* = 12.5, 5.1 Hz, 1H, H6_pro-R,biotin_), 2.57 (d, *J* = 12.5 Hz, 1H, H6_pro-S,biotin_), 2.39 (d, *J* = 1.3 Hz, 3H, 4_AMC_-CH_3_), 2.03 (t, *J* = 7.5 Hz, 2H, NHCOC*H*_*2*_), 1.85 (s, 3H, CH_3_CO), 1.79–1.69 (m, 1H, Hβ_lys,A_), 1.69–1.55 (m, 3H, Hβ_lys,B_, Hγ_leu,_C*H*_*2,A*_C4_biotin_), 1.53–1.21 (m, 11H, Hβ_leu_,Hγ_lys_,Hδ_lys_,NHCOCH_2_C*H*_*2*_C*H*_*2*_C*H*_*2,B*_C4_biotin_), 0.88 (d, *J* = 6.6 Hz, 3H, CH_3,leu,A_), 0.84 (d, *J* = 6.5 Hz, 3H, CH_3,leu,B_). ^13^C NMR (DMSO-*d*_6_) *δ* 172.9 (CO_leu_), 171.9 (CONHε_lys_), 171.4 (CO_lys_), 169.7 (CO_ac_), 169.0 (CO_gly_), 162.8 (C2_biotin_), 160.0 (C2_AMC_), 153.6 (C8a_AMC_), 153.1 (C4_AMC_), 142.1 (C7_AMC_), 125.9 (C5_AMC_), 115.3 (C6_AMC_), 115.1 (C4a_AMC_), 112.4 (C3_AMC_), 105.8 (C8_AMC_), 61.1 (C3a_biotin_), 59.2 (C6a_biotin_), 55.4 (C4_biotin_), 53.6 (Cα_lys_), 51.5 (Cα_leu_), 42.1 (Cα_gly_), 40.5 (Cβ_leu_), 39.8 (C6_biotin_), 38.2 (Cε_lys_), 35.2 (NHCO*C*H_2_), 31.4 (Cβ_lys_), 28.9 (Cδ_lys_), 28.2 (NHCOCH_2_CH_2_*C*H_2_), 28.0 (*C*H_2_C4_biotin_), 25.3 (NHCOCH_2_*C*H_2_), 24.2 (Cγ_leu_), 22.9 (CH_3,leu,A_), 22.8 (Cγ_lys_), 22.5 (*C*H_3_CO), 21.6 (CH_3,leu,B_), 18.0 (4_AMC_–CH_3_). HRMS *m*/*z* calcd for C_36_H_52_N_7_O_8_S^+^ [M+H]^+^, 742.3593; found, 742.3605.

#### Ac-Glu-Thr-Asp-Lys(isobutyryl)-AMC (6f)

The title compound was synthesized according to the general acylation procedure. Reagents: isobutyric acid (5 mg, 0.06 mmol, 1.2 equiv), HATU (23 mg, 0.06 mmol, 1.3 equiv), lutidine (12 mg, 0.12 mmol, 2.6 equiv). Reaction time: 4 h. Purification of the crude residue by preparative HPLC afforded Ac-Glu-Thr-Asp-Lys(isobutyryl)-AMC (**6f**, 17 mg, 47%) as a colorless fluffy solid after lyophilization. ^1^H NMR (600 MHz, DMSO-*d*_6_) δ 10.22 (s, 1H, NH_AMC_), 8.22 (d, *J* = 7.7, 1H, NH_asp_), 8.12 (d, *J* = 7.7, 1H, NH_glu_), 7.98 (d, *J* = 7.5, 1H, NH_lys_), 7.78 (d, *J* = 2.0, 1H, H8_AMC_), 7.71 (d, *J* = 8.7, 1H, H5_AMC_), 7.68 (d, *J* = 8.0, 1H, NH_thr_), 7.65 (t, *J* = 5.6, 1H, NHε_lys_), 7.52 (dd, *J* = 8.7, 2.0, 1H, H6_AMC_), 6.26 (d, *J* = 1.2, 1H, H3_AMC_), 4.61 (m, 1H, Hα_asp_), 4.34–4.28 (m, 2H, Hα_glu_, Hα_lys_), 4.25 (dd, *J* = 8.0, 4.4, 1H, Hα_thr_), 4.05–3.99 (m, 1H, Hβ_thr_), 3.01 (m, 2H, Hε_lys_), 2.75 (dd, *J* = 16.7, 5.7, 1H, Hβ_asp-A_), 2.59 (dd, *J* = 16.7, 7.4, 1H, Hβ_asp-B_), 2.40 (d, *J* = 1.1, 3H, CH_3-AMC_), 2.33–2.24 (m, 3H, Hγ_glu_, NHCOC*H*(CH_3_)_2_), 1.96–1.89 (m, 1H, Hβ_glu-A_), 1.86 (s, 3H, H_ac_), 1.78–1.70 (m, 2H, Hβ_glu-B_, Hβ_lys-A_), 1.67–1.60 (m, 1H, Hβ_lys-B_), 1.43–1.22 (m, 4H, Hγ_lysA-B_, Hδ_lys_), 1.05 (d, *J* = 6.3, 3H, Hγ_thr_), 0.95 (dd, *J* = 6.8, 3.0, 6H, NHCOCH(C*H*_3_)_2_). ^13^C NMR (151 MHz, DMSO) δ 175.9 (CONHε_lys_), 174.0 (COδ_glu_), 171.9 (CO_asp_), 171.6 (COα_glu_), 171.1 (CO_lys_), 170.6 (CO_asp_), 169.9 (CO_thr_), 169.6 (CO_ac_), 160.0 (CO_AMC_), 153.6 (C8a_AMC_), 153.1 (C4_AMC_), 142.0 (C7_AMC_), 125.9 (C5_AMC_), 115.4 (C6_AMC_), 115.1 (C4a_AMC_), 112.3 (C3_AMC_), 105.8 (C8_AMC_), 66.7 (Cβ_thr_), 57.9 (Cα_thr_), 53.9 (Cα_lys_), 52.1 (Cα_glu_), 49.6 (Cα_asp_), 38.2 (Cε_lys_), 35.8 (Cβ_asp_), 34.0 (NHCO*C*H(CH_3_)_2_), 31.3 (Cβ_lys_), 30.2 (Cγ_glu_), 28.8 (Cδ_lys_), 27.0 (Cβ_glu_), 22.8 (Cγ_lys_), 22.4 (C_ac_), 19.6 (NHCOCH(*C*H_3_)_2_), 19.2 (Cγ_thr_), 18.0 (CH_3-AMC_). HRMS *m*/*z* calcd for C_35_H_48_N_6_NaO_13_^+^ [M+Na]^+^, 783.3177; found, 783.3184.

#### Ac-Glu-Thr-Asp-Lys(2-methylacrylyl)-AMC (**6j**)

The title compound was synthesized according to the general acylation procedure. Reagents: methacrylic acid (5 mg, 0.06 mmol, 1.2 equiv), HATU (23 mg, 0.06 mmol, 1.3 equiv), lutidine (12 mg, 0.12 mmol, 2.6 equiv). Reaction time: 6 h. Purification of the crude residue by preparative HPLC afforded Ac-Glu-Thr-Asp-Lys(2-methylacrylyl)-AMC (**6j**, 26 mg, 74%) as a colorless fluffy solid after lyophilization. ^1^H NMR (600 MHz, DMSO-*d*_6_) δ 12.22 (bs, 2H, COOH_asp_, COOH_glu_), 10.21 (s, 1H, NH_AMC_), 8.22 (d, *J* = 7.7 Hz, 1H, NH_asp_), 8.12 (d, *J* = 7.7 Hz, 1H, NH_glu_), 8.00 (d, *J* = 7.5 Hz, 1H, NH_lys_), 7.86 (t, *J* = 5.7 Hz, 1H, NHε_lys_), 7.77 (d, *J* = 2.0 Hz, 1H, H8_AMC_), 7.71 (d, *J* = 8.7 Hz, 1H, H5_AMC_), 7.68 (d, *J* = 8.0 Hz, 1H, NH_thr_), 7.51 (dd, *J* = 8.7, 2.0 Hz, 1H, H6_AMC_), 6.26 (d, *J* = 1.2 Hz, 1H, H3_AMC_), 5.60 (s, 1H, NHCOC(CH_3_)C*H*_2_), 5.27 (m, 1H, NHCOC(CH_3_)C*H*_2_), 4.61 (m, 1H, Hα_asp_), 4.32 (m, 2H, Hα_glu_, Hα_lys_), 4.25 (dd, *J* = 8.0, 4.4 Hz, 1H, Hα_thr_), 4.05–3.98 (m, 1H, Hβ_thr_), 3.09 (m, 2H, Hε_lys_), 2.75 (dd, *J* = 16.7, 5.7 Hz, 1H, Hβ_asp-A_), 2.59 (dd, *J* = 16.7, 7.5 Hz, 1H, Hβ_asp-B_), 2.39 (s, 3H, CH_3-AMC_), 2.31–2.23 (m, 2H, Hγ_glu_), 1.97–1.89 (m, 1H, Hβ_glu-A_), 1.86 (s, 3H, H_ac_), 1.82 (m, 3H, NHCOC(C*H*_3_)CH_2_), 1.75 (m, 2H, Hβ_glu-B_, Hβ_lys-A_), 1.65 (m, 1H, Hβ_lys-B_), 1.45 (m, 2H, Hδ_lys_), 1.41–1.32 (m, 1H, Hγ_lys-A_), 1.27 (m, 1H, Hγ_lys-B_), 1.05 (d, *J* = 6.3 Hz, 3H, Hγ_thr_). ^13^C NMR (151 MHz, DMSO) δ 174.0 (COδ_glu_), 171.9 (COγ_asp_), 171.6 (CO_glu_), 171.2 (CO_lys_), 170.7 (CO_asp_), 169.9 (CO_thr_), 169.6 (CO_ac_), 167.4 (COε_lys_), 160.0 (CO_AMC_), 153.6 (C8a_AMC_), 153.1 (C4_AMC_), 142.0 (C7_AMC_), 140.1 (NHCO*C*(CH_3_)CH_2_), 125.9 (C5_AMC_), 118.7 (NHCOC(CH_3_)*C*H_2_), 115.3 (C6_AMC_), 115.2 (C4a_AMC_), 112.4 (C3_AMC_), 105.8 (C8_AMC_), 66.7 (Cβ_thr_), 57.9 (Cα_thr_), 53.9 (Cα_lys_), 52.2 (Cα_glu_), 49.6 (Cα_asp_), 38.7 (Cε_lys_), 35.8 (Cβ_asp_), 31.3 (Cβ_lys_), 30.3 (Cγ_glu_), 28.8 (Cδ_lys_), 27.0 (Cβ_glu_), 22.9 (Cγ_lys_), 22.4 (C_ac_), 19.2 (Cγ_thr_), 18.7 (NHCOC(*C*H_3_)CH_2_), 18.0 (CH_3-AMC_). HRMS *m*/*z* calcd for C_35_H_46_N_6_NaO_13_^+^ [M+Na]^+^, 781.3021; found, 781.3025.

#### Ac-Glu-Thr-Asp-Lys(2-methylcrotonoyl)-AMC (6k)

The title compound was synthesized according to general acylation procedure. Reagents: 2-methylcrotonic acid (6 mg, 0.06 mmol, 1.2 equiv), HATU (23 mg, 0.06 mmol, 1.3 equiv), lutidine (12 mg, 0.12 mmol, 2.6 equiv). Reaction time: overnight. Purification of the crude residue by preparative HPLC afforded Ac-Glu-Thr-Asp-Lys(2-methylcrotonoyl)-AMC (**6k**, 21 mg, 60%) as a colorless fluffy solid after lyophilization. ^1^H NMR (600 MHz, DMSO-*d*_6_) δ 10.21 (s, 1H), 8.22 (d, *J* = 7.7 Hz, 1H, NH_asp_), 8.12 (d, *J* = 7.7 Hz, 1H, NH_glu_), 7.99 (d, *J* = 7.5 Hz, 1H, NH_lys_), 7.77 (d, *J* = 2.0 Hz, 1H, H8_AMC_), 7.71 (d, *J* = 8.7 Hz, 1H, H5_AMC_), 7.68 (m, 2H, NHε_lys_), 7.51 (dd, *J* = 8.7, 2.0 Hz, 1H, H6_AMC_), 6.28–6.23 (m, 2H, COC(CH_3_)C*H*CH_3_, H3_AMC_), 4.61 (m, 1H, Hα_asp_), 4.35–4.28 (m, 2H, Hα_glu_, Hα_lys_), 4.25 (dd, *J* = 8.0, 4.4 Hz, 1H, Hα_thr_), 4.05–3.99 (m, 1H, Hβ_thr_), 3.08 (m, 2H, Hε_lys_), 2.75 (dd, *J* = 16.7, 5.7 Hz, 1H, Hβ_asp-A_), 2.59 (dd, *J* = 16.7, 7.5 Hz, 1H, Hβ_asp-B_), 2.40 (d, *J* = 1.2 Hz, 3H, CH_3 AMC_), 2.30–2.23 (m, 2H, Hβ_glu_), 1.96–1.88 (m, 1H, Hα_glu-A_), 1.86 (s, 3H, H_ac_), 1.78–1.71 (m, 2H, Hα_lys-A_, Hα_glu-B_), 1.71–1.69 (m, 3H, COC(C*H*_3_)CHCH_3_), 1.68–1.61 (m, 4H, COC(CH_3_)CHC*H*_3_, Hα_lys-B_), 1.47–1.40 (m, 2H, Hδ_lys_), 1.39–1.31 (m, 1H, Hγ_lys-A_), 1.31–1.22 (m, 1H, Hγ_lys-B_), 1.05 (d, *J* = 6.3 Hz, 3H, Hγ_thr_). ^13^C NMR (151 MHz, DMSO) δ 174.0 (COδ_glu_), 171.9 (COγ_asp_), 171.6 (CO_glu_), 171.2 (CO_lys_), 170.7 (CO_asp_), 169.9 (CO_thr_), 169.6 (CO_ac_), 168.3 (COε_lys_), 160.0 (CO_AMC_), 153.6 (C8a_AMC_), 153.1 (C4_AMC_), 142.0 (C7_AMC_), 132.0 (CO*C*(CH_3_)CHCH_3_), 128.8 (COC(CH_3_)*C*HCH_3_), 125.9 (C5_AMC_), 115.3 (C6_AMC_), 115.1 (C4a_AMC_), 112.3 (C3_AMC_), 105.8 (C8_AMC_), 66.7 (Cβ_thr_), 57.9 (Cα_thr_), 53.9 (Cα_lys_), 52.2 (Cα_glu_), 49.6 (Cα_asp_), 38.7 (Cε_lys_), 35.8 (Cβ_asp_), 31.3 (Cβ_lys_), 30.2 (Cβ_glu_), 28.8 (Cδ_lys_), 27.0 (Cα_glu_), 22.8 (Cγ_lys_), 22.4 (C_ac_), 19.2 (Cγ_thr_), 18.0 (CH_3 AMC_), 13.5 (COC(CH_3_)CH*C*H_3_), 12.3 (COC(*C*H_3_)CHCH_3_). HRMS *m*/*z* calcd for C_36_H_48_N_6_NaO_13_^+^ [M+Na]^+^, 795,3177; found, 795.3184.

#### Ac-Glu-Thr-Asp-Lys(3-methylcrotonoyl)-AMC (6l)

The title compound was synthesized according to general acylation procedure. Reagents: 3-methylcrotonic acid (6 mg, 0.06 mmol, 1.2 equiv), HATU (23 mg, 0.06 mmol, 1.3 equiv), lutidine (12 mg, 0.12 mmol, 2.6 equiv). Reaction time: after 3 h, methylcrotonic acid (1 mg, 0.01 mmol, 0.2 equiv), HATU (4 mg, 0.01 mmol, 0.2 equiv), and lutidine (2 mg, 0.01 mmol, 0.3 equiv) were added to the reaction mixture. The reaction mixture was then stirred overnight. Purification of the crude residue by preparative HPLC afforded Ac-Glu-Thr-Asp-Lys(3-methylcrotonoyl)-AMC (**6l**, 23 mg, 63%) as a colorless fluffy solid after lyophilization. ^1^H NMR (600 MHz, DMSO-*d*_6_) δ 10.22 (s, 1H, NH_AMC_), 8.23 (d, *J* = 7.7, 1H, NH_asp_), 8.12 (d, *J* = 7.7, 1H, NH_glu_), 7.99 (d, *J* = 7.5, 1H, NH_lys_), 7.77 (d, *J* = 2.0, 1H, H8_AMC_), 7.71 (d, *J* = 8.7, 1H, H5_AMC_), 7.70–7.66 (m, 2H, NHε_lys_, NH_thr_), 7.52 (dd, *J* = 8.7, 2.0, 1H, H6_AMC_), 6.26 (d, *J* = 1.2, 1H, H3_AMC_), 5.60 (d, *J* = 1.3, 1H, NHCOC*H*C(CH_3_)_2_), 4.62 (m, 1H, Hα_asp_), 4.36–4.28 (m, 2H, Hα_lys_, Hα_glu_), 4.26 (dd, *J* = 8.0, 4.3, 1H, Hα_thr_), 4.05–3.99 (m, 1H, Hβ_thr_), 3.04 (m, 2H, Hε_lys_), 2.75 (dd, *J* = 16.7, 5.7, 1H, Hβ_asp-A_), 2.59 (dd, *J* = 16.7, 7.4, 1H, Hβ_asp-B_), 2.40 (d, *J* = 1.2, 3H, CH_3-AMC_), 2.31–2.23 (m, 2H, Hγ_glu_), 2.04 (d, *J* = 1.0, 3H, NHCOCHC(C*H*_3_)_2_), 1.96–1.89 (m, 1H, Hβ_glu-A_), 1.86 (s, 3H, H_ac_), 1.79–1.70 (m, 5H, NHCOCHC(C*H*_3_)_2_, Hβ_lys-A_, Hβ_glu-B_), 1.68–1.60 (m, 1H, Hβ_lys-B_), 1.44–1.32 (m, 3H, Hδ_lys_, Hγ_lys-A_), 1.32–1.22 (m, 1H, Hγ_lys-B_), 1.05 (d, *J* = 6.3, 3H, Hγ_thr_). ^13^C NMR (151 MHz, DMSO) δ 174.0 (COδ_glu_), 171.9 (COγ_asp_), 171.6 (CO_glu_), 171.1 (CO_lys_), 170.7 (CO_asp_), 169.9 (CO_thr_), 169.6 (CO_ac_), 165.9 (COε_lys_), 160.0 (CO_AMC_), 153.6 (C8a_AMC_), 153.1 (C4_AMC_), 147.9 (NHCOCH*C*(CH_3_)_2_), 142.0 (C7_AMC_), 125.9 (C5_AMC_), 119.2 (NHCO*C*HC(CH_3_)_2_), 115.4 (C6_AMC_), 115.2 (C4a_AMC_), 112.4 (C3_AMC_), 105.8 (C8_AMC_), 66.7 (Cβ_thr_), 57.9 (Cα_thr_), 53.9 (Cα_lys_), 52.2 (Cα_glu_), 49.6 (Cα_asp_), 38.0 (Cε_lys_), 35.8 (Cβ_asp_), 31.3 (Cβ_lys_), 30.3 (Cγ_glu_), 28.9 (Cδ_lys_), 27.0 (Cβ_glu_), 26.7 (NHCOCHC(*C*H_3_)_2_), 22.9 (Cγ_lys_), 22.4 (C_ac_), 19.2 (NHCOCHC(*C*H_3_)_2_, Cγ_thr_), 18.0 (C_CH3-AMC_). HRMS *m*/*z* calcd for C_36_H_48_N_6_NaO_13_^+^ [M+Na]^+^, 795.3177; found, 795.3182.

